# TEX13B is important for germ cell development and male fertility

**DOI:** 10.1101/2022.01.11.475851

**Authors:** Umesh Kumar, Digumarthi V S Sudhakar, Nithyapriya Kumar, Hanuman T Kale, Rajan Kumar Jha, Nalini J Gupta, B N Chakravarthy, Mamata Deenadayal, Aarti Deenadayal Tolani, Swasti Raychaudhuri, P Chandra Shekar, Kumarasamy Thangaraj

## Abstract

The recent epidemiological studies suggest that nearly one out of every 7 reproductive age couples face problem to conceive a child after trying for at least one year. Impaired fertility of the male partner is causative in approximately 50% of the infertile couples. However, the etiologies of large proportion of male infertility are still unclear. Our unpublished exome sequencing data identified several novel genes including TEX13B, which motivated us to further explore the role of TEX13B in male infertility in large infertile case control cohort. Hence in this study, we have examined the role of *TEX13B* in male infertility by whole gene sequencing 628 infertile and 427 control men and have demonstrated the functional role of Tex13b in spermatogonia GC1spg (GC1) cells. We identified 2 variants on TEX13B which are tightly associated with male infertility. TEX13B gene exclusively expressed in germ cells, but its molecular functions in germ cells are still unknown. Hence, we demonstrated the functional importance of Tex13b in GC1 cell line by genomic manipulation via CRISPR-Cas9 and mass spectrometry-based whole cell proteomics. The gene knock out in GC1 cell line clearly shows that Tex13b play an important role in germ cell growth and morphology. We demonstrate that Tex13b knockout or conditional overexpression in GC1 cells reprograms the metabolic status from an oxidative phosphorylation to glycolysis state and vice versa. In conclusion, our study clearly showed the importance of Tex13b in germ cells development and Its association with male infertility.

## Introduction

Infertility, defined as inability of a couple to conceive a child after trying to conceive for one year is a major reproductive health problem in about 15 percent reproductive age couples around the globe (ref). Impaired fertility of the male partner is causative risk factor in approximately 50% of the infertile couples. Data from our laboratory has shown that about 8.5% infertility among Indian men is due to the Y chromosome microdeletion. Further, analysis of several autosomal genes (*CAMK4*, *UBE2B* and *TNP2-4*) accounted for additional 17.5% genetic factors responsible for infertility among Indian men (ref). However, etiologies of large proportion of male infertility are still unclear therefore, it is essential to identify the novel causative mutations for male infertility.

Recent genomic studies revealed that the mutations in mammalian X chromosome have direct impact on fertility because X chromosome is enriched for genes involved in spermatogenesis (Stouffs et al., 2009). A recent study has shown that an x chromosome gene *TEX11* mutations were a common cause of meiotic arrest and azoospermia in infertile men (Yang et al., 2015). Another study has stress on the X chromosome genes which express exclusively male germ cells, identified 10 X-linked genes, including Tex13b in mouse as well as human homolog of Tex13, *TEX13B*. *TEX13B* is orthologous to mouse gene *Tex13b* that is found to be is involved in transcriptional regulation during spermatogenesis (Wang et al., 2001). Furthermore, recently it has been shown that Tex13b express specifically pre-leptotene stage of the spermatogonia cells, indicating its potent role in spermatogenesis. The importance of Tex13b in spermatogonia differentiation is indicated in another study since Tex13b found to be most highly connected to the genes specific to germ cells by hub-gene-network analysis (Liao et al., 2017). However, the molecular function *TEX13B* gene is yet to be explored in germ cells and its association in male infertility.

Since TEX13B has been found as a novel hit in our exome sequencing (unpublished), we have sequenced the coding region (exon 2 and 3) of *TEX13B* in 628 infertile men (nature of infertile patients) along with 427 ethnically matched fertile controls. We have identified two variants in the coding region of *TEX13B* and found to be significantly associated with azoospermia patients. To explore the molecular function of TEX13B we created the Tex13b knockout spermatogonia GC1spg (GC1) cells. We specifically choose GC1 cells to characterized Tex13b because these cells belong to pre-leptotene stage of germ cells (spermatogonia B) and Tex13 expresses specifically at this stage (Hofmann, Narisawa, Hess, & Millan, 1992; Wang et al., 2001). Since Tex13b shown to be a transcription factor, hence, to examine the differential protein expression in Tex13b knockout cells we performed isotope labeling by amino acid in cell culture (SILAC) and mass spectrometry-based proteomics. Results in this study clearly show that Tex13b regulate balance between OXPHOS and glycolysis.

## Results

### TEX13B variants associated with male infertility

*TEX13B* (Testis-Expressed Protein 13B) is another candidate gene identified by exome sequencing with the non-synonymous variant rs41300872 (p.Gly197Arg) showing significant association with male infertility (OR=1.77, 95% CI); *P*=0.002. *TEX13B* is a gene located on X-chromosome and is exclusively expressed in male germ cells and spermatogonia (Wang, McCarrey, Yang, & Page, 2001). To identify additional genetic variants, we sequenced the complete coding region of *TEX13B* in 628 infertile men (443 NOA, 105 OAT and 80 severe oligozoospermia individuals) along with 427 ethnically matched fertile control men (**Table 1)**. We found an additional rare variant, rs775429506 (p.Gly237Glu) exclusively in two NOA men.

**Table 1:** List of samples collected from different centers in India for *TEX13B* sequencing Each PCR was carried out in 10.0 μl reaction mixture using 2X EmeraldAmp GT PCR master mix® (Takara Bio INC, Japan), 1.0 pM of each primer. Primer sequences and PCR conditions are shown in **Table 2**.

| Source of sample<br>(Total number of cases) | Infertile men |  |  | Fertile men |
| --- | --- | --- | --- | --- |
|  | Azoospermia | Severe OA | OAT |  |
| IIRC, Hyderabad (250) | 115 | 80 | 55 | 141 |
| IRM, Kolkata (378) | 328 | 0 | 50 | 286 |
| Total | 443 | 80 | 105 | 427 |
|  | 628 |  |  |  |

**Table 2:**
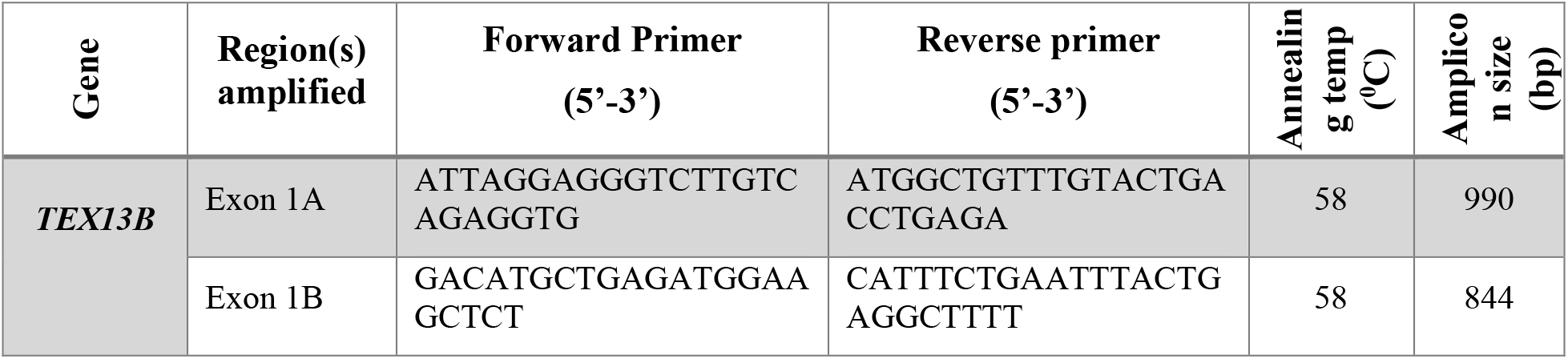
Primer sequences and PCR conditions used to amplify TEX13B gene

### Tex13b regulates germ cell proliferation, mitochondrial OXPHOS

To understand the role of *TEX13B* in spermatogenesis, we performed functional studies using mouse-derived GC-1spg (GC1 cells) cells. GC1 cells belong to the spermatogonia B stage and *Tex13b* (mouse homolog of *TEX13B*) is specifically expressed at this stage (Hofmann, Narisawa, Hess, & Millan, 1992; Wang et al., 2001). We deleted the first coding exon of *Tex13b* in GC1 cells using the CRISPR-Cas9 **(Figure 1A, Supplementary figure 1A-1C)**and observed morphological differences (multipolar cells with large extensions), and reduced cell proliferation **(Figure 1B, Figure 1C)**. Since the molecular function of Tex13b protein is unknown, we examined the differential protein expression in knockout cells (GTKO31) compared to GC1 cells using mass spectrometry of whole-cell isotope-labeled proteins (**Figure 1D**). A total of 145 (out of 3278) proteins showed significantly altered expression in GTKO31 cells compared to wildtype cells **(Supplementary file 1)**. Results were validated by qPCR of their respective mRNAs of top fourteen differentially expressed proteins in two independent *Tex13b* knockout clones (GTKO7 and GTKO31) compared to wildtype GC1 cells **(supplementary figure 1D and 1E)**. Interestingly, the gene ontology (GO) of most of the downregulated proteins in GTKO31 belong to mainly two protein complexes, the mitochondrial electron transport chain protein complexes (ETCP) and the mitochondrial ribosomes (mitoribosome) complex. **(Figure 1E and 1F, Supplementary file 1)**. These results were further validated by immunostaining of a ETCP complex I protein NDUFB4 (**Figure 1G and 1H**) and immunoblotting of multiple ETCP complex I proteins **(Figure 1I, Figure 1J)**.

**Figure 1:**
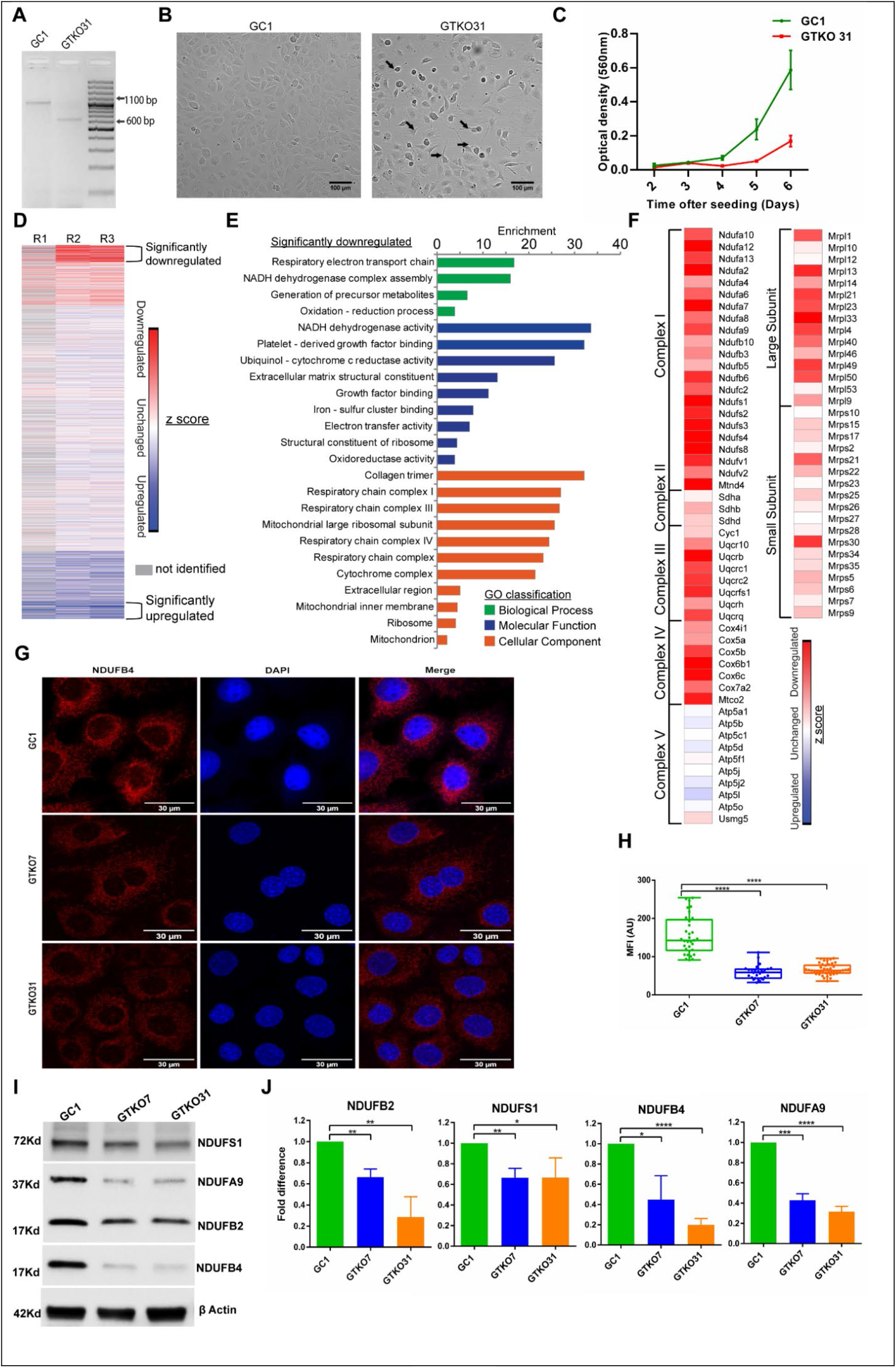
Functional characterization of *TEX13b* in GC1 cell lines. A) PCR based genotyping of *Tex13b* locus of GTKO31 cells compared to GC1 cells; B) Microscopic analysis of phonotypic changes in GTKO31 cells compared to GC1. The black arrows show the multipolar cells with multiple extensions in GTKO31 cells; C) Impact of the *Tex13b* deletion in GTKO31 cells on cell proliferation compared to wild type GC1 cells; D) Heat map of z-score for H/L (heavy isotope/low isotope) intensity of identified peptides in mass spectrometry in three independent replicates R1, R2, R3; E) Gene ontology of significantly down-regulated proteins in GTKO31 cells; F) The heat map of the Z-score of the identified mitochondrial ETCP proteins and mitochondrial ribosomal proteins in GTKO31 cells. G) Immunostaining of NDUFB4 in GC1, GTKO7 and GTKO31 cells; H) The mean fluorescence intensity of NDUFB4 in GTKO7 and GTKO31 cells compared to GC1 cells; I) Immunoblotting of multiple ETCP Complex I proteins in *Tex13b* knockout clones GTKO7 and GTKO31 cells compared to GC1 cells; J) Western blot quantification of multiple ETCP proteins in GTKO7 and GTKO31 compared to wild-type GC1 cells. The data is represented as mean +/− SD in biological triplicates.

Further, to understand the effect of ETCP down regulation in on cellular respiration we performed respirometry analysis and observed that all respiration parameters were significantly repressed in GTKO31 cells compared to wild type GC1 cells **(Figure 2A)**. There are two possibilities of these observations, either there is reduction of total abundance of mitochondria in *Tex13b* knockout cells or there is specific down regulation of ETCP while unchanged mitochondrial abundance. To check these possibilities, we performed mitochondrial staining and flow cytometric analysis (**Figure 2B**). We found that there is no significant change in total mitochondrial abundance in *Tex13b* knockout GTKO31 cells compared to GC1 cells (**Figure 2C**). Further, the proteomics results indicate that there is no significant change in protein expression of Complex V of ETCP (**Figure 1F**). Altogether these results clearly suggest that *Tex13b* knockout leads to specific downregulation of ETCP complexes, mainly Complex I, Complex II, Complex III and Complex IV which leads to depression of OXPHOS.

**Figure 2:**
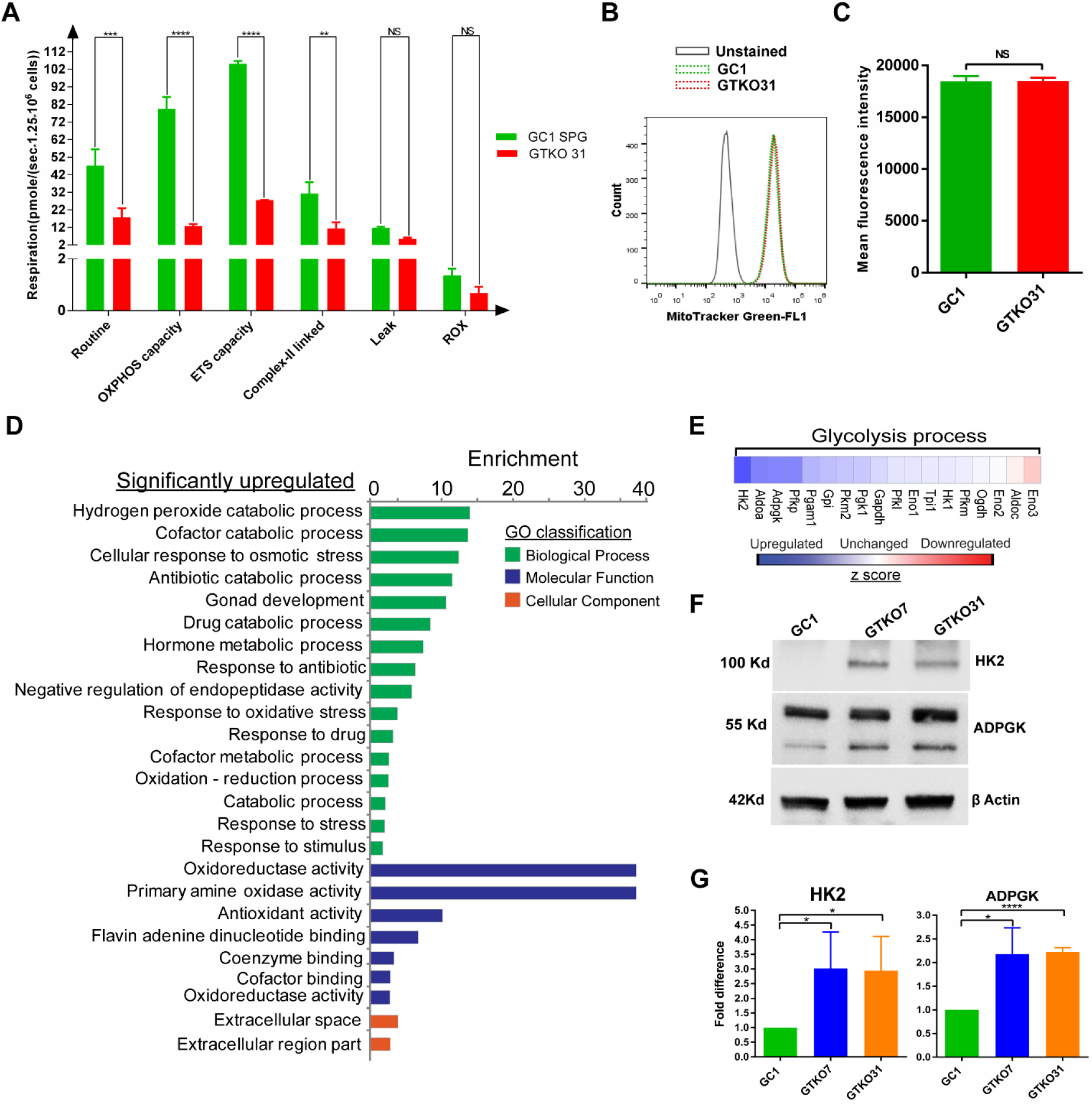
Repressed repression and upregulated genes in *Tex13b* knockout GC1 cells-. A) Differential mitochondrial respiration parameters in GTKO31 cells compared to GC1 cells.; B) Flowcytometric analysis of MitoTracker™ green stained GC1 and GTKO31 cells. C) Mean fluorescence intensity of MitoTracker™ green fluorescence of GTKO31 compared to GC1 cells in three independent flowcytometric experiments; D) GO analysis of the upregulated proteins in GTKO31 cells; E) The heat map of the Z-score of the identified glycolytic enzymes in proteomics experiment; F) Immunoblotting of glycolytic enzyme HK2 and ADPGK2 in *Tex13b* knockout clones GTKO7 and GTKO31 cells compared to GC1 cells G) Western blot quantification of glycolytic enzymes HK2 and ADPGK in GTKO7 and GTO31 compared to GC1 cells. The data is represented as mean +/− SD in biological triplicate.

The gene ontology of the upregulated proteins in *Tex13b* knockout cells belongs several different cellular processes including gonadal development (**Figure 2D**). Interestingly, one of the gene ontology processes is related to catabolism (**Figure 2D**). Glycolysis and OXPHOS are two interdependent energy sources for cell survival. Since OXPHOS is repressed in *Tex13b* knockout cells we examine the expression level of glycolytic enzymes from proteomics data. We found that the expression of most of the identified glycolytic enzymes are upregulated in *Tex13b* knockout cells compare to the GC1 cells (**Figure 2E**). These results were further validated by immunostaining of two important glycolytic enzymes, hexokinase 2 (HK2) and ADP dependent glucokinase (ADPGK) (**Figure 2F and 2G**).

### Tex13b overexpression leads to upregulation of ETCP proteins

To understand role of *Tex13b* further, in regulation of metabolism, we created a Doxycycline-inducible Flag-*Tex13b* overexpressing GC1 cell line GTIG (**Figure 3A, Figure 3B**). we observed that doxycycline induced *Tex13b* overexpression leads to significant increase in the abundance of multiple ETCP proteins **(Figure 3C, Figure 3D)**, whereas there is no significant increase of ETCP protein in doxycycline-treated wild-type GC1 cells (**Figure 3E, Figure 3F**). These results shows that *Tex13b* regulates the abundance of mitochondrial proteins and positively regulates the ETCP proteins. The results altogether indicate that loss of function of Tex13b leads to shift in energy metabolism from OXPHOS to glycolysis, known as Warburg effect or aerobic glycolysis (Warburg, Wind, & Negelein, 1927).

**Figure 3:**
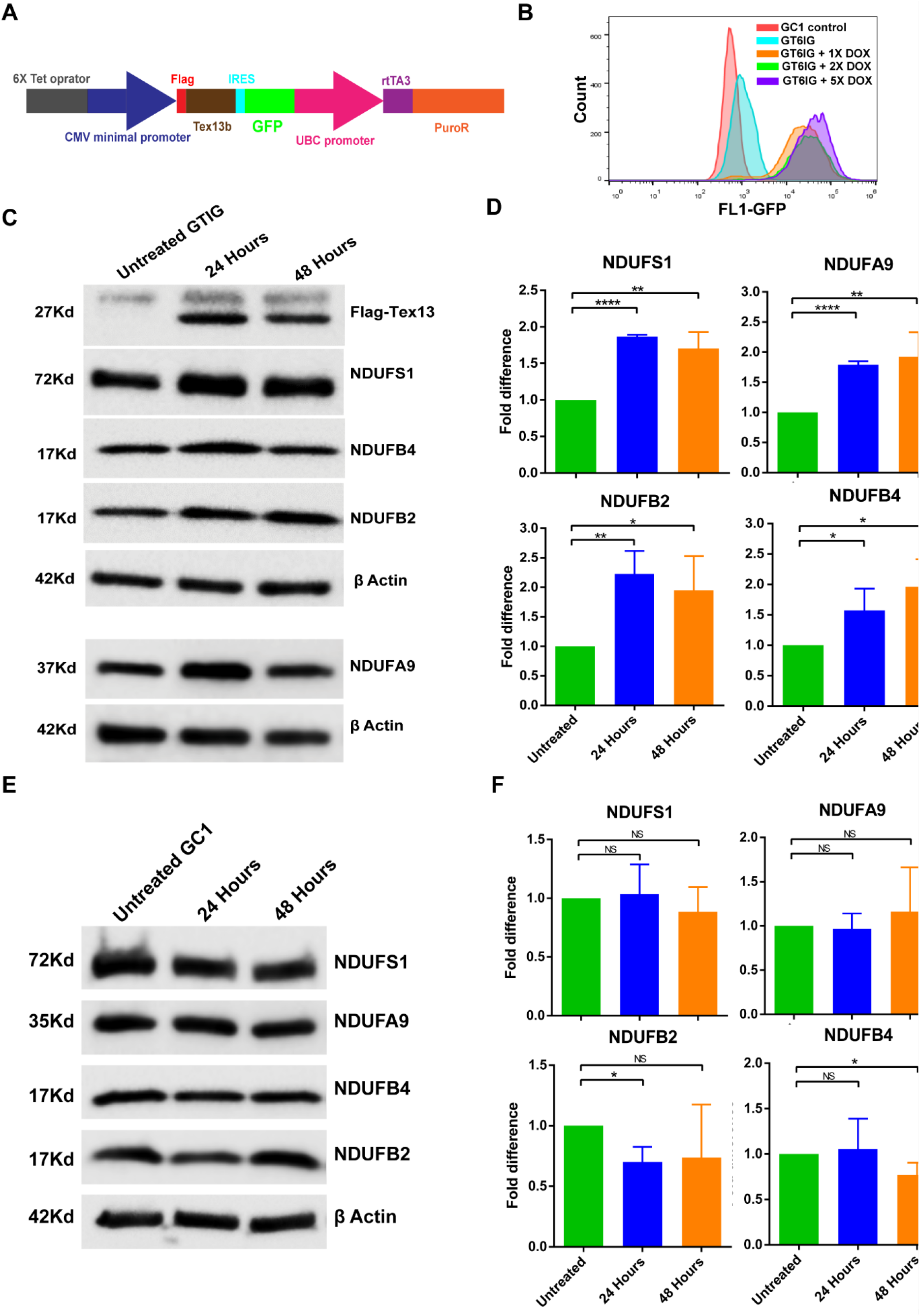
Conditional over-expression of *Tex13b* in GC1 cells leads to upregulation of ETCP. A) Schematic depiction of doxycycline inducible Flag-Tex13b-IRES-GFP vector construct; B) The flowcytometric analysis of the doxycycline inducible GC1 Flag-Tex13b-IRES-GFP (GTIG) positive clone C) Western blot analysis of ETCP proteins and quantification (D) of doxycycline induced Flag-Tex13b overexpressing GTIG cells compared to mock induced GTIG cells at two independent time points. E) Western blot analysis and quantification (F) of ETCP proteins of doxycycline treated wild type GC1 cells compared to mock treated GC1 cells in three independent experiments

## Discussion

*TEX13B*, is an X-chromosomal gene that belongs to *TEX* gene family and is exclusively expressed in male germ cells (Bellil, Ghieh, Hermel, Mandon-Pepin, & Vialard, 2021; Wang et al., 2001). In mouse, the *Tex13* gene family has four members namely *Tex13a*, *Tex13b*, *Tex13c1*, and *Tex13d* (Kwon et al., 2016). Mouse *Tex13* genes are transcribed specifically or predominantly in male germ cells, and their expression is developmentally regulated. Wang et al., identified two human orthologues of *TEX13*-*TEX13A* and *TEX13B* and demonstrated their exclusive expression in testis (Wang et al., 2001). Recent studies have shown that changes in energy metabolism and interplay of mitochondrial OXPHOS and glycolysis is a fundamental event during spermatogenesis and play an important role in cell fate determination during the primordial germ cell specification, pluripotent stem cells differentiation and induced pluripotent stem cell reprograming (Guo et al., 2017; Hayashi et al., 2017). Hence, the regulation of the switching from glycolysis to OXPHOS and vice versa could be an important factor for germ cell lineage specification and fate determination. Here, we propose that *Tex13b* disruption in GC1 cells leads to a shift in their metabolism from OXPHOS to glycolysis. Since Tex13b is transcription factor and localized to the nucleus, it might regulate the nuclear coded mitochondrial OXPHOS genes, transcriptionally, either directly or indirectly by regulating the transcription of OXPOS genes (Figure 4). Thus, *Tex13b* might be important for germ cell differentiation and maintenance of cell potency by regulating the balance between OXPHOS and glycolysis. Thus, our results clearly demonstrate that *Tex13b* is important for GC1 cells growth and proliferation. The genetic perturbation of *Tex13b* leads to a shift in energy metabolism pathways *in vitro*.

**Figure 4:**
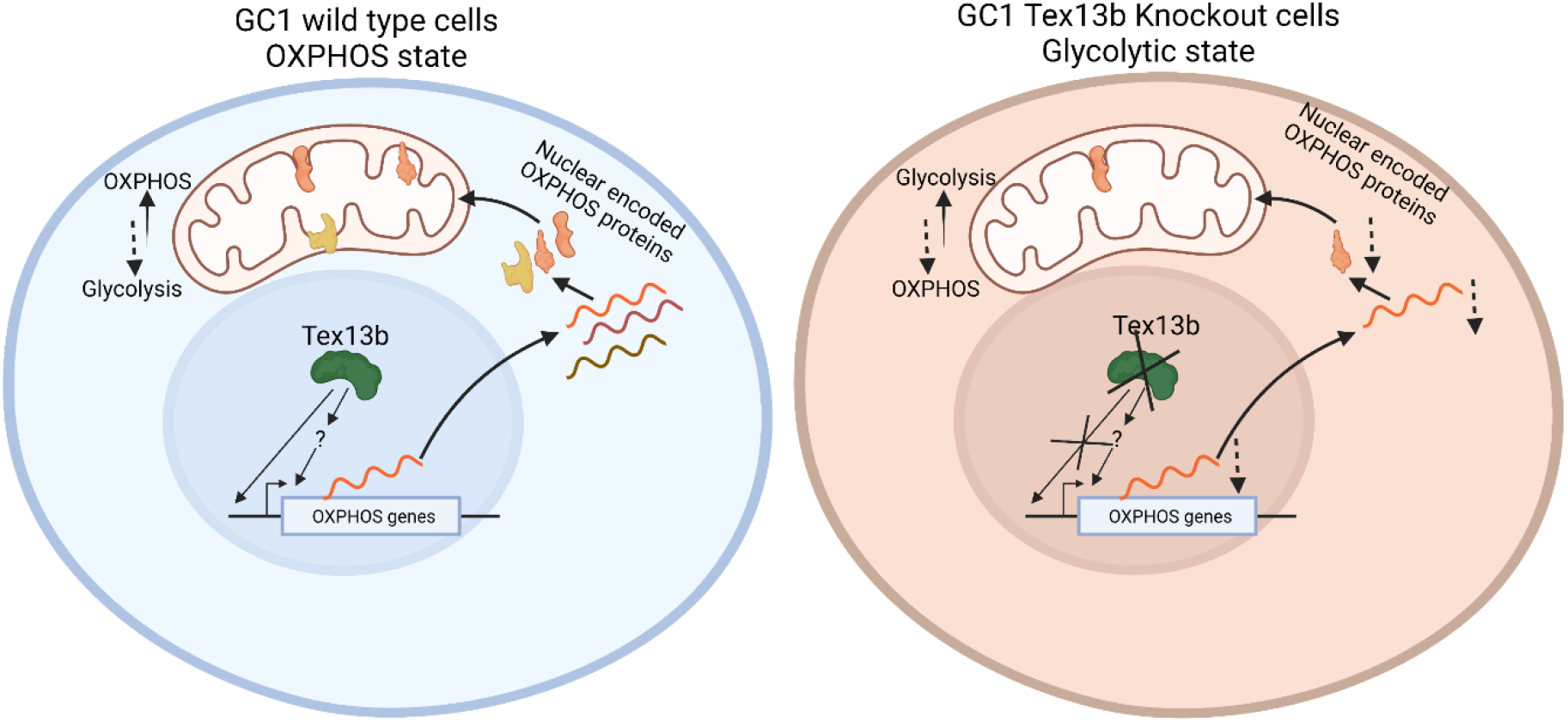
Functional model for Tex13b mediated regulation of metabolism. In the wildtype cells Tex13b express and localized to nucleous. Tex13b leads to positively regulate the nuclear encoded OXPHOS genes, and leads to a normal state where GC1 cells depends upon OXPHOS for energy metabolism. Whereas in Tex13b knockout cells the transcriptional activation of nuclear encoded OXPHOS genes is alterd due to ubsence of Tex13b. Consequently, the nuclear encoded OXPHOS proteins are down regulated in Tex13b knockout cells, which leads to upregulation of glycolytic enzymes. The dashed arrows indicating the down regulation of OXPHOS proteins.

## Materials and methods

### Subjects

Selection of infertile men was done after careful clinical evaluation by expert urologists. Infertile men were classified based on semen parameters, strictly following the WHO criteria (WHO manual, 5th edition, 2010). Individuals with known cause of infertility such as; varicocele, chromosomal abnormalities, testicular trauma, obstructions in the sperm delivery system and endocrine abnormalities were excluded from this study. About 5.0 ml of peripheral blood sample was collected from each individual after obtaining their informed written consent. Genomic DNA was isolated from the blood samples using phenol-chloroform method described in our earlier study(Thangaraj et al., 2003). All the infertile men were screened for Y chromosome microdeletions using STS markers that are specific to *AZF*a, *AZF*b, *AZF*c regions(Thangaraj et al., 2003). Infertile men, who did not possess any Y chromosome microdeletion and or mutation in known autosomal genes were included in the present study.

### *TEX13B* gene sequencing

A total of 628 infertile men (443 NOA, 105 OAT and 80 severe oligozoospermic individuals) and 427 ethnically matched fertile control men were included in this study. Of these, 378 idiopathic infertile men (328 azoospermia and 50 severe Oligoasthenoteratozoospermia) and 286 fertile men were recruited from the Institute of Reproductive Medicine (IRM), Kolkata, India and 250 infertile men (115 azoospermia and 55 Oligoasthenoteratozoospermia and 80 severe oligozoospermia) and 141 fertile controls were recruited from Infertility Institute and Research Centre (IIRC), Hyderabad, India shown in (**Table 1**).

### Statistical analysis

To estimate the risk or association of alleles with infertility, allele and genotypic frequencies were compared between the case and control groups using Statistical Package for Social Sciences, ver.16 (SPSS Inc., USA) or PLINK 1.9. In a χ2 test for association and Odds ratios (OR) at their 95% confidence intervals (95% CI) were calculated and p-values less than 0.05 were considered statistically significant.

### *Tex13b* Knock-out

The GC1 cells were maintained in DMEM media containing 10% FBS. To create genetic deletion in *Tex13b,* we constructed a Cas9-twin guide RNA plasmid with high fidelity Cas9 (spCas9-HF1) tagged with green fluorescence protein (GFP) using self-cleaving peptide T2A linker, regulated by chicken actin promoter and oligos encoding two guide RNAs (CACCGCTCAGCCATCATGAATTGCG and CACCGGAAATAAGACTGGGTTAGG) in single construct regulated by PU6 promoters as depicted in **Supplementary figure 1A**. The first guide RNA was targeted for 5^’^UTR of *Tex13b* and start codon junction and the second guide RNA for second intron-exon junction. 10 μg of Cas9 twin guide construct was nucleofected in GC1 cells using 4D-Nucleofector (LONZA, Switzerland) and after growing them for 24 hours, the GFP positive cells (Cas9 positive) were sorted by FACS. The sorted cells were grown in single cell colony for 15 days and PCR based genotyping was done to screen clones for genetic deletion using forward primer (5’-AAGGGCGAGCTAAAGAACTGATGG-3’) and reverse primer (5’-CATGATTGGCATCTGTGCAACTGG-3’). The overall strategy has been shown in **Supplementary figure 1B.** The precise deletion was confirmed by sequencing of the *Tex13b* targeted locus. Two independent clones (GC1 *Tex13b* Knock-out) were named as GTKO7 and GTKO31.

### *Tex13b* conditional over-expression

To study the gain of function of *Tex13b* in GC1 cells, we created a doxycycline inducible Flag-Tex13b expressing GC1 cell line, named as GTIG (GC1 Tex13b IRES-GFP). The plasmid was designed and the construct with Flag-tag *Tex13b*, regulated with 6X Tet operator, whereas the rtTA 3 (binds to Tet operator only in presence of doxycycline) tagged with puromycin resistant protein was regulated with ubiquitin promoter as depicted in the **Figure 3A**. The *Tex13b* was also tagged with *GFP* in fusion with internal ribosomal entry site (IRES) sequence. The 10 μg of plasmid construct was nucleofected into the GC1 cells and the integration positive cells were selected by growing them in presence of 2 μg/ml puromycin in growth media for 7 days. The selected cells were serially diluted to create single cell colony and screened by flowcytometric analysis of GFP expression upon 24 hours of doxycycline treatment **(Figure 3B)**. The *Tex13b* induction were further confirmed by western blotting of Flag-Tex13b by Flag specific antibody **(Figure 3C)**.

### Mass spectroscopy and proteomics analysis

To explore the possible targets and function of *Tex13b* gene in germ cells, we examined the global differential abundance of proteins in *Tex13b* KO GC1 cells (GTKO31) compared to wild type GC1 cells. We labelled whole cell proteins by growing the cells in Stable Isotope Labelling of Amino acids in Cell culture (SILAC), DMEM media (Thermo Fisher Scientific) supplemented with 10% dialysed serum and with heavy isotopes L-lysine 2HCl (^13^C_6_, ^15^N_2_)/L-arginine HCl (^13^C_6_, ^15^N_4_) (Lys8/Arg10)] (GC1 cells) or light isotope [L-lysine 2HCl/L-arginine HCl (Lys0/Arg0)] (GTKO31 cells) for 6 passages. The labelling efficiency was checked using mass spectrometry. The equal number of labelled GC1 and GTKO31 cells were mixed, and protein was extracted using RIPA buffer (50 mM Tris HCl, 150 mM NaCl, 1.0% (v/v) NP-40, 0.5% (w/v) Sodium Deoxycholate, 1.0 mM EDTA, 0.1% (w/v) SDS and protease inhibitor cocktail). 100 μg protein from mixed GTKO31 and GC1 mixed cells were separated using gradient acrylamide gel (Invitrogen®) and stained by Coomassie Brilliant Blue (CBB). The gel was de-stained overnight, and stained proteins was cut into nine different fractions according to size and abundance of bands. These gel fractions were processed, and proteins was digested into peptide with in-gel tryptic digestion method according to protocol(Shevchenko, Wilm, Vorm, & Mann, 1996). The peptide was extracted and desalted using C18 columns using previously described protocol(Rappsilber, Ishihama, & Mann, 2003). The peptides were dissolved in 2% formic acid and analysed on a Q-Exactive HF mass spectroscopy (Thermo Fisher Scientific). The peptides were separated using an EASY-Spray PepMap RSLC C18 column at flow rate of 300 nL/minute on a 60 min linear gradient of the mobile phase (5% acetonitrile (ACN) containing 0.2% formic acid (buffer A) and 95% ACN containing 0.2% formic acid (buffer B)).

The peptide was identified from the acquired raw mass data using MaxQuant (Ver. 1.3.0.5) and searched in *Mus musculus* dataset of the Swiss-Prot data base(Cox & Mann, 2008). For every identified peptide, the H/L (intensity of heavy labelled peptide/ intensity of light labelled peptide) ratios were calculated and were converted into log2 space. The mean ratios and standard deviations were calculated for each data peptide for three independent experiments. The log2H/L ratio of each protein was converted into a z-score as described in the previous study(Rawat, Ghosh, Mondal, Anusha, & Raychaudhuri, 2020). Differing confidence level cut-offs were applied to the z-score analysis to determine proteins that were significantly differentially regulated with 95 % confidence level, corresponding to z-scores more or less than ±1.960. Using a 95% cut off, significant differential regulation was observed for 158 proteins in GTKO cells (66 upregulated, 79 downregulated).

### Real-time quantitative PCR

The mRNA concentration (of selected genes) was compared by real time PCR. The RNA was isolated and was reverse transcribed into cDNA using cDNA synthesis Kit (Takara). The quantitative PCR was done using power SYBR Green PCR master mix (Invitrogen). The sequence of the primers, used in real time PCR is listed in **Table 3.**GAPDH was used as reference gene for normalization.

**Table 3:**
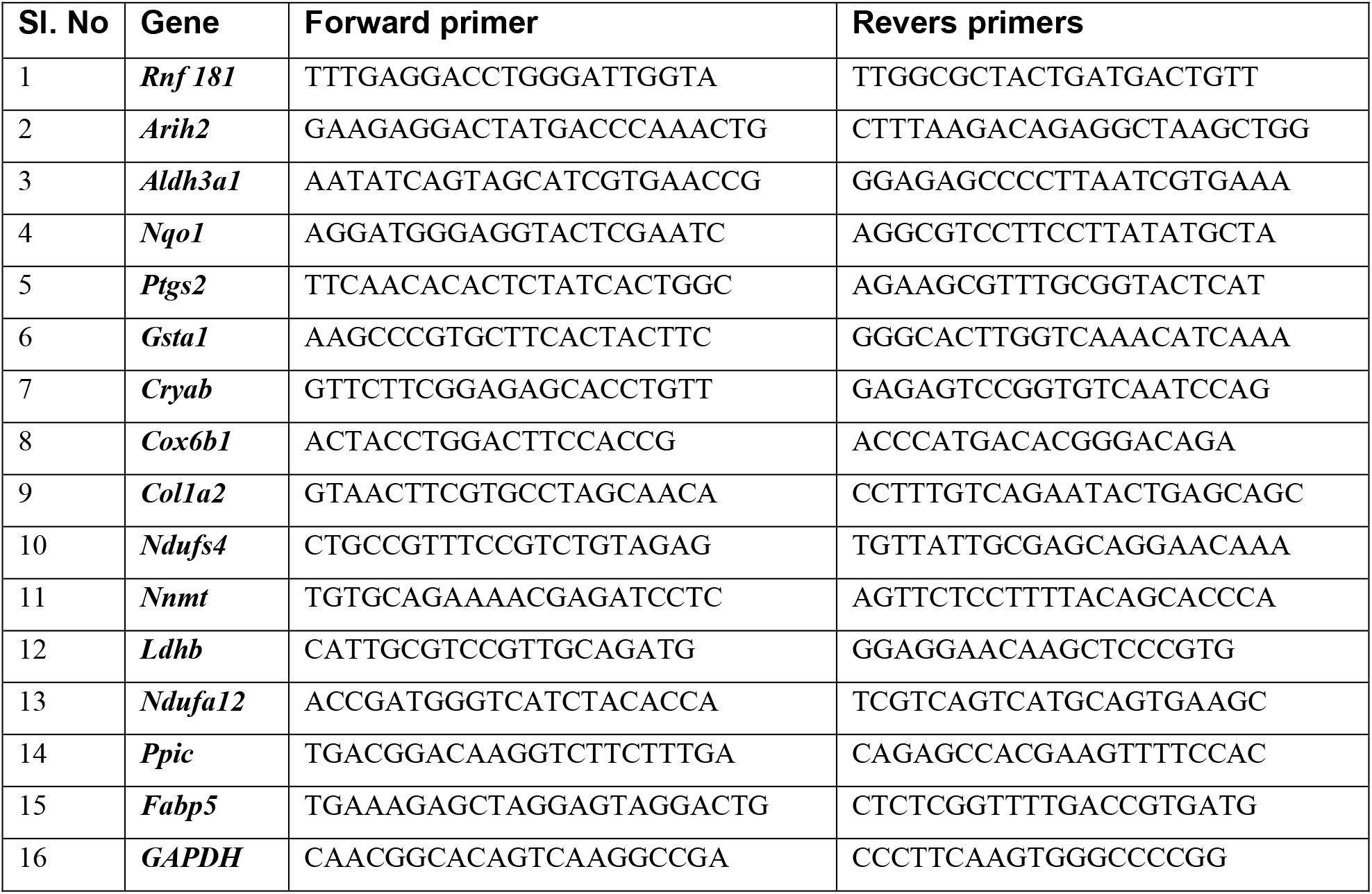
Primers used for qPCR to validate targets identified from mass spectrometry data

### Mitochondrial respirometric analysis

To know the mitochondrial OXPHOS status GTKO31 cells in comparison with GC1 cells, the respirometric measurements were performed using High Resolution Respirometer (O2K, OROBOROS® Instrument, Innsbruck, Austria) equipped with DatLab7 recording and analysis software. The oxygraph measurements were performed using 1.25*10^6^ cells each GC1 and GTKO separately in Mitochondrial Respiration Medium (Mir05) (Oroboros® Instruments, Innsbruck, Austria) at 37°C with continuous stirring. The substrate-uncoupler-inhibitor-titration **(SUIT)**protocol was applied to both control GC1 and knockout GTKO31 cell lines in a closed chamber according to standard protocol (Makrecka-Kuka, Krumschnabel, & Gnaiger, 2015; Pesta et al., 2011). Different parameters of respirations were calculated for GTKO31 in comparison with GC1 cells and plotted.

### Mitochondrial abundance quantification

To study mitochondrial abundance GTKO31 cells in comparison with GC1 cells we stained the cells with MitoTracker green. About 2 lack cells was diluted into staining media and incubated for 30 minutes in CO_2_ incubator at 37°C temperature. The cells were centrifuged and diluted in PBS and green fluorescence was analysed by flow cytometer.

### Proliferation assay

To evaluate the role of *Tex13b* in germ cells growth we did MTT assay for GTKO31 cells in compare with GC1 cells. The MTT assay was done every 24 hours according to protocol described in the previous study(Kumar & Kumar, 2015). Briefly, 500 cells were seeded for each GC1 and GTKO31 cells per well (12 wells each) of 96 well plate in DMEM growth media in 6 different plates and media was changed every 48 hrs. Every 24-hour media from one plate was removed and cells were incubated with media containing 5 μM MTT for 2 h at 37°C. After that the media was removed and purple crystals which was formed during incubation was dissolved in 100 microliter of dimethyl sulfoxide (DMSO) and absorbance was recorded at 562 nm. The proliferation was calculated by comparing the absorbance of GC1 cells compare to GTKO31 cells.

### Immunoblotting

The cell lysis was performed using RIPA lysis buffer at 4°C temperature for 30 minutes. The lysate was further sonicated in cell rupture. The protein concentration was estimated and fractionated on polyacrylamide gel electrophoresis and transferred on to PVDF membrane. Different proteins were detected using respective antibodies (**Table 4)**.

**Table 4:** List of Antibodies used in immunoblotting and immunostaining

| Sl.No. | Antibodies | Company (Catalogue) |
| --- | --- | --- |
| 1) | Anti-NDUFS1 antibody | abcam (ab169540) |
| 2) | Anti-NDUFB4 antibody | abcam (ab192243) |
| 3) | Anti-NDUFA9 antibody | abcam (ab14713) |
| 4) | Anti-NDUFB2 antibody | abcam (ab186748) |
| 5) | Anti-Hexokinase II antibody | Abcam (ab209847) |
| 6) | Anti-ADPGK antibody | Abcam (ab228633) |
| 7) | Anti-β-Actin | Thermo Fisher Scientific |

## Acknowledgements

We thank Shivali Rawat for her assistance in proteomics data analysis, Anurupa Moitra for her assistance in *TEX13B* cell biology work. KT acknowledge the Council of Scientific and Industrial Research (CSIR) for funding under the network project (BSC0101 and MLP0113). KT thank SERB, Department of Science and Technology, Government of India for the J C Bose Fellowship.

---

**Supplement Figure 1:**
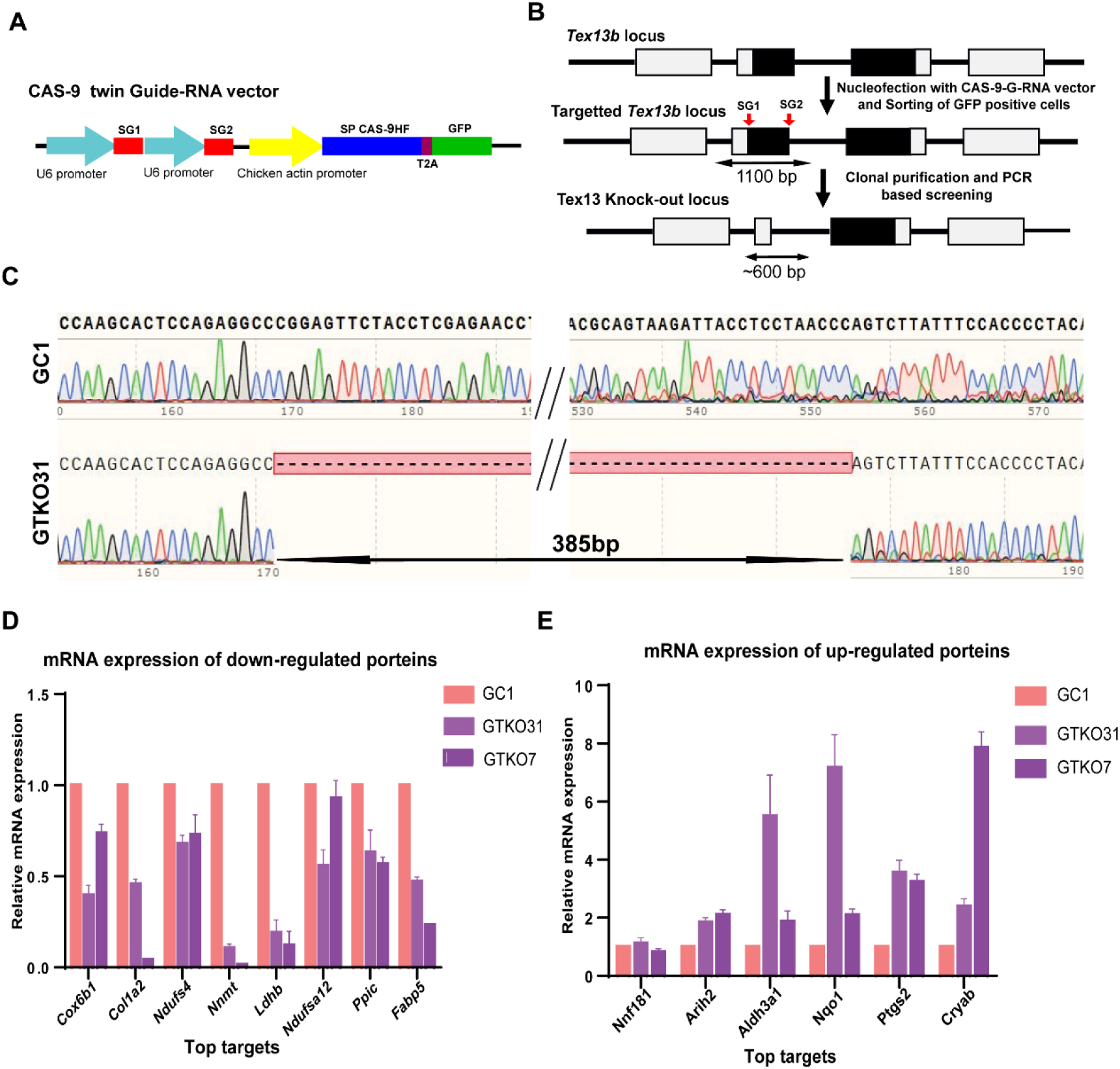
*Tex13b* knockout in GC1 cells. A) Schematic depiction of CRISPR-Cas9 twin guide-RNA vector construct. SG1 and SG2 are two guides, specific for the *Tex13b* gene B) Schematic representation of the strategy for the creation of *Tex13b* knockout GC1 cells. PCR based screening for positive clones was done using forward (For) and reverse (Rev) primers; C) Electropherogram showing *Tex13b* locus in GTKO31 cells compared to the wild-type GC1 cells; D) qPCR analysis of the top 8 downregulated proteins and E) top 6 upregulated proteins in proteomics experiment in two independent *Tex13b* knockout clones compared to the wild-type GC1 cells.

